# Early suppression of excitability in subcortical band heterotopia modifies epileptogenesis in rats

**DOI:** 10.1101/2022.09.06.506850

**Authors:** Delphine Hardy, Emmanuelle Buhler, Dmitrii Suchkov, Antonin Vinck, Aurélien Fortoul, Françoise Watrin, Alfonso Represa, Marat Minlebaev, Jean-Bernard Manent

## Abstract

Malformations of cortical development represent a major cause of epilepsy in childhood. However, the pathological substrate and dynamic changes leading to the development and progression of epilepsy remain unclear. Here, we characterized an etiology-relevant rat model of subcortical band heterotopia (SBH), a diffuse type of cortical malformation associated with drug-resistant seizures in humans. We used longitudinal electrographic recordings to monitor the age-dependent evolution of epileptiform discharges during the course of epileptogenesis in this model. We found both quantitative and qualitative age-related changes in seizures properties and patterns, accompanying a gradual progression towards a fully developed seizure pattern seen in adulthood. We also dissected the relative contribution of the band heterotopia and the overlying cortex to the development and age-dependent progression of epilepsy using timed and spatially targeted manipulation of neuronal excitability. We found that an early suppression of neuronal excitability in SBH slows down epileptogenesis in juvenile rats, whereas epileptogenesis is paradoxically exacerbated when excitability is suppressed in the overlying cortex. However, in rats with active epilepsy, similar manipulations of excitability have no effect on chronic spontaneous seizures. Together, our data support the notion that complex developmental alterations occurring in both the SBH and the overlying cortex concur to creating pathogenic circuits prone to generate seizures. Our study also suggests that early and targeted interventions could potentially influence the course of these altered developmental trajectories, and favorably modify epileptogenesis in malformations of cortical development.

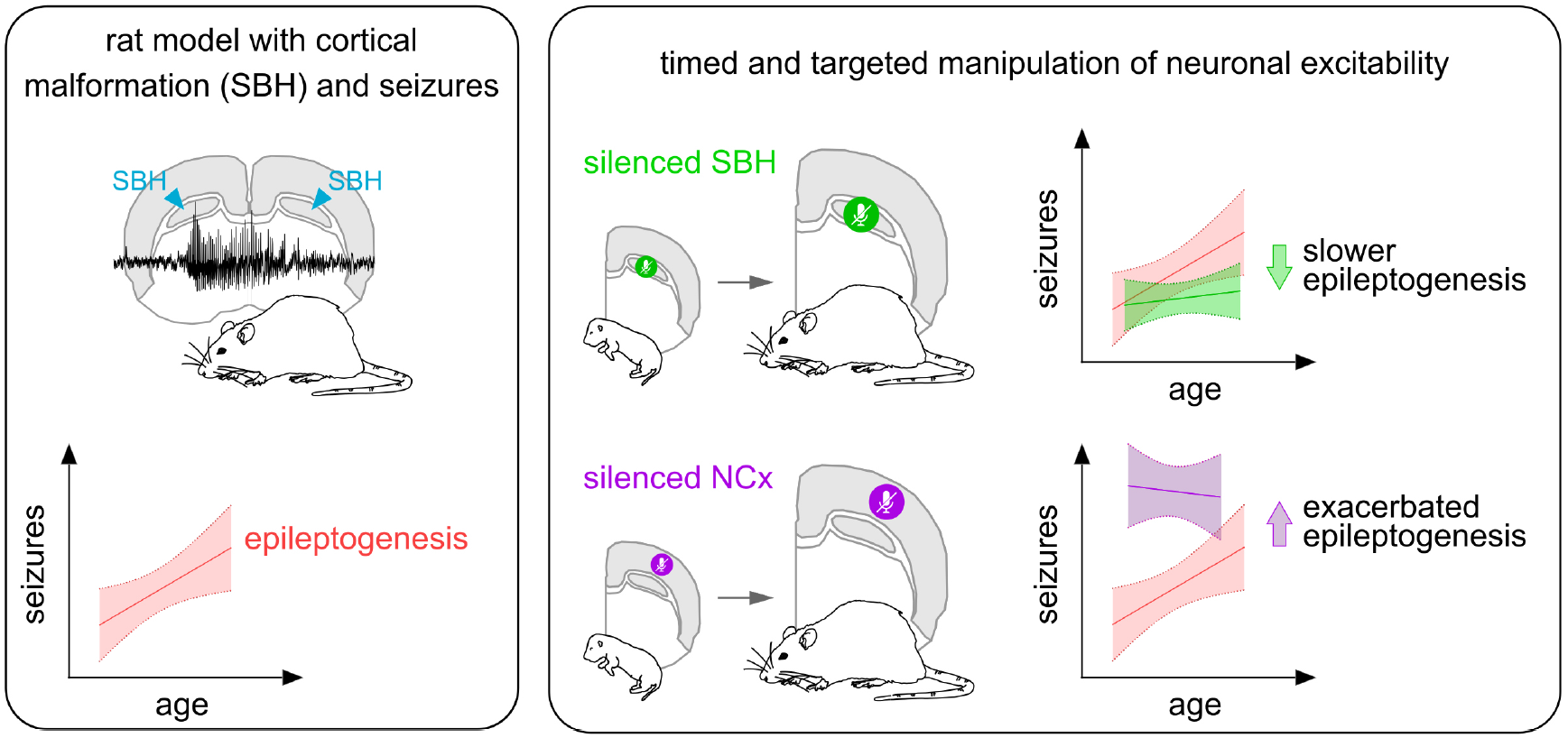

## Introduction

Epileptogenesis refers to the development and extension of tissue capable of generating spontaneous seizures, resulting in the development of an epileptic condition and/or the progression of the epilepsy after it is established.^1^ Various acquired or genetic factors can trigger epileptogenesis, and, accordingly, there is a wide spectrum of epilepsy syndromes, each of them characterized by distinct etiologies, natural course of disease and underlying pathological mechanisms. Identifying the dynamic changes occurring during the course of epileptogenesis in order to prevent, ameliorate, or cure epilepsy is a major challenge for epilepsy research.^2^

Disrupted formation of the cerebral cortex represents a major cause of epilepsy in childhood, and malformations of cortical development (MCDs) account for up to 40% of intractable or drug-resistant cases.^3^ MCDs comprise a large group of structural brain abnormalities, with genetic and non-genetic underlying causes, and variable clinical presentations and burden of disability. One can broadly distinguish two main groups: early-onset and diffuse MCDs, generally associated with poor neurological outcomes, and late-onset, more focal MCDs, associated with milder outcomes.^4^ Because MCDs are usually not detected until patients begin having epileptic seizures, our understanding of epileptogenesis in MCDs remains limited.

Subcortical band heterotopia (SBH) belongs to the group of diffuse MCDs,^5^ and is characterized by a band of grey matter separated from the cortex and lateral ventricles by zones of white matter.^6,7^ DCX mutations are the most frequently diagnosed genetic causes for SBH, accounting for 100% of familial cases and over 80% of sporadic cases.^8,9^ Patients with SBH present with mild to moderate intellectual disability (68% of cases), seizures (85-96%), including a high proportion of drug-resistance (78%).^9,10^ Although our understanding of the pathophysiology of epileptogenic MCDs, has greatly improved,^11^ both the epileptogenic substrate and dynamic changes occurring during the course of epileptogenesis remain largely elusive, especially in severe and diffuse cases of MCDs such as SBH.

Here, we characterized the age-dependent evolution of electrographic discharges during epileptogenesis in an etiology-relevant rat model of SBH^12^ using longitudinal electrocorticographic monitoring. We found that epilepsy gradually progresses with age, with both quantitative and qualitative changes of seizure properties and patterns. We also dissected the relative contribution of the SBH and the overlying cortex to the development and age-dependent progression of epilepsy using a targeted approach for suppressing neuronal excitability. We report that an early suppression of neuronal excitability in SBH slows down epileptogenesis in juvenile rats, whereas epileptogenesis is paradoxically exacerbated when excitability is suppressed in the overlying cortex. However, in rats with active epilepsy, similar manipulations of excitability have no effect on chronic spontaneous seizures. Together, these findings suggest that complex developmental alterations occurring in both the SBH and the overlying cortex concur to creating pathogenic circuits prone to generate seizures. Our study also suggest that early and targeted interventions could potentially influence the course of these altered developmental trajectories, and favorably modify epileptogenesis.

## Materials and methods

### Animal ethics

Animal experiments were performed in agreement with European directive 2010/63/UE and received approval [2020080610441911_v2(APAFIS#26835)] from the French Ministry for Research after ethical evaluation by the Institutional Animal Care and Use Committee of Aix-Marseille University.

### Tripolar in utero electroporation

We performed tripolar in utero electroporation as described.^12,13^ Timed pregnant Wistar rats (Janvier) received buprenorphine (Buprecare, 0.03mg/kg) and Carprofen (Rimadyl, 5mg/kg) and were anesthetized with isoflurane (2.5%) 30 min later. Uterine horns were exposed under isoflurane anesthesia, and plasmid vectors (plasmid conditions, Supplementary Fig. 1 and 5) were microinjected bilaterally into the lateral ventricles of embryonic day 16 embryos, together with fast green dye. Tripolar electroporations were accomplished by delivering 50V voltage pulses (BTX ECM830 electroporator; Harvard Apparatus) across tweezer-type electrodes laterally pinching the head of each embryo through the uterus, and a third electrode positioned at the brain midline (Supplementary Fig. 1A). Successfully electroporated rats were selected one day after birth (P1) based on fluorescent protein expression by using transcranial illumination.

### Telemetric electrocorticographic recording

We performed telemetric recordings of electrocorticographic signals (ECoG) in freely behaving, unrestrained Dcx-KD rats as described.^12^ Juvenile bilateral Dcx-KD rats received buprenorphine (Buprecare, 0.03mg/kg) and Carprofen (Rymadyl, 5mg/kg) and were anesthetized with isoflurane (induction 5%, maintenance 2-3%) 30 min later. Rats were placed in a stereotaxic frame (Kopf Instruments) and 2 holes were drilled through the skull above somatosensory cortex for electrode implantation using coordinates relative to Bregma (AP:-0.5, L:+3.5). Reference electrode was placed on the cerebellum (AP:-10, L:+1.4). Electrodes were secured with screws and mounted with dental acrylic. A subcutaneous pocket was created to insert the telemetry transmitter (CTA-F40, Data Sciences International) on the left dorsal side of the rat. ECoG signals were digitized with Ponemah software (recording schedule as in Fig. 2).

### Seizure detection

ECoG signals were processed as follows (Supplementary Fig. 2): 1) we extracted 150 ms segments every 10 ms, 2) we normalized amplitudes and extracted mean values from the normalized signal, 3) we calculated velocities, 4) we generated 50 by 50 matrices and filled them with the paired values of amplitude/velocities calculated for every ECoG segments. 5) We vectorized the resulting images depicting the phase portraits of individual segments to generate the input dataset.

Because we found that the phase portraits of SWs were distinct to that of rhythmic waves or background noise (Fig. 1E2, 1F), we used their characteristic shapes as alphabet characters to develop a seizure detection framework based on a character recognition approach. We used a neural network with 112 hidden layers and cross entropy parameter for the network performance estimation. The training dataset was composed of 38 911 events (34 485 noise events, 2 162 rhythmic waves and 2 264 SWs) that we manually selected using threshold methods and visual inspection. After training, the trained neural network was then used to detect seizures on the target dataset generated from ECoG signals processed similarly (Supplementary Fig. 2B).

**Figure 1.**
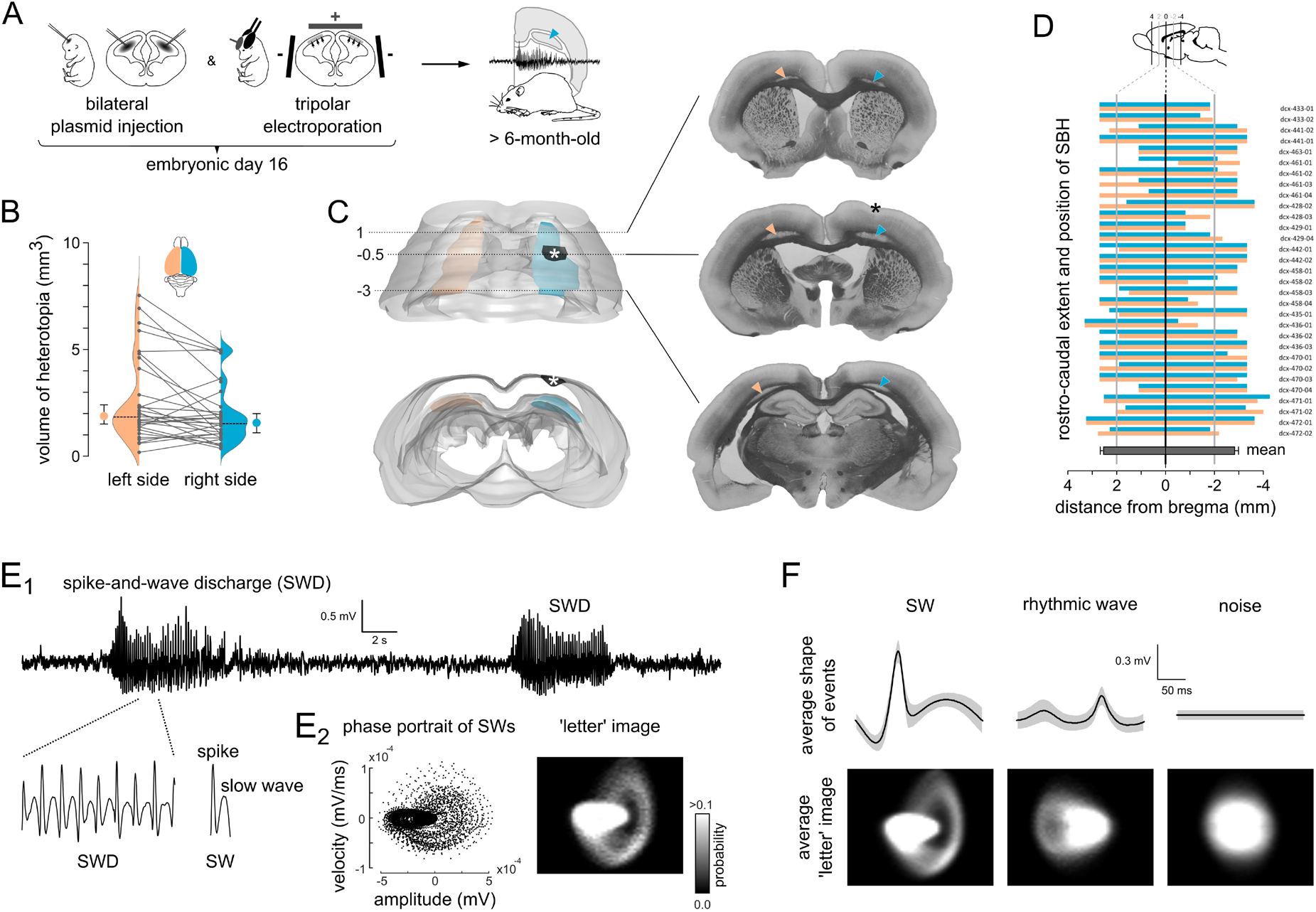
Adult bilateral Dcx-KD rats harbor bilateral SBH and chronic spontaneous seizures. **(A)** Experimental timeline. **(B)** Line graphs and density estimates showing the volume of SBH in the left (orange) and right (blue) hemispheres of 31 adult bilateral Dcx-KD rats. Grey dots correspond to individual values, with the left and right values for individual rats connected with a line. Dashed lines on the density plots show medians. Orange and blue circles with error bars show medians and 95% confidence intervals. **(C)** Top view (top) and rear view (bottom) of the rostral part of a three-dimensionally reconstructed adult brain with bilateral SBH (in orange in the left hemisphere, in blue in the right hemisphere). (right) 3 frontal sections taken at the indicated positions from bregma showing the band heterotopia (orange and blue arrowheads) in the white matter (in dark grey). Asterisks indicate the electrode implantation site. **(D)** Clustered bar graphs showing the rostrocaudal extent and position of SBH relative to bregma in the 31 bilateral Dcx-KD rats analyzed, in the left (orange) and right (blue) hemispheres. Dark grey bar shows the mean values with SEM. Animal identifier codes are given next to each pairs of bars. **(E1)** Representative ECoG trace showing two spike-and-wave discharges (SWD), composed of trains of spikes (S) followed by associated slow waves (W), and referred to as spike-and-wave (SW) complexes. **(E2)** Phase portrait of 1,000 SWs (left) and corresponding ‘letter’ image. **(F)** Average traces of SW, rhythmic wave and noise, and corresponding ‘letter’ image of their phase portraits.

### Seizure analysis

We defined a seizure episode as a sequence of SW complexes with inter-event intervals shorter than 500 ms. If intervals were longer than 500 ms, we considered that a new seizure episode has started. Only seizures with durations greater than 1 s were used for further analysis. For every detected seizures, we extracted peak values as follows: 1) We detrended seizures from their low frequency components (less 1 Hz); 2) We smoothed seizures with a 10 ms window; 3) We extracted phase components using Hilbert transformation; 4) We identified points with rapid changes in phase, which corresponds to peaks position. For each peak, we calculated time-stamp, amplitude and half-width.

### Slice electrophysiology

P13 to P16 rats were deeply anesthetized with tiletamine/zolazepam (Zoletil, 40 mg/kg) and medetomidine (Domitor, 0.6 mg/kg), and decapitated. The brain was then quickly removed and was placed in chilled and oxygenated ACSF containing the following (in mM): 25 NaHCO3, 1.25 NaH2PO4.H2O, 6.3 D-glucose, 2.5 KCl, 7 MgCl2.6H2O, 0.5 CaCl2.2H2O and 132.5 choline chloride. Coronal slices (250-μm-thick) were obtained using a vibrating microtome (Leica Biosystems). During the electrophysiological experiments, the slices were superfused with oxygenated ACSF at a rate of 2 ml/min containing the following (in mM): 126 NaCl, 26 NaHCO3, 1.2 NaH2PO4.H2O, 6.3 D-Glucose, 3.5 KCl, 1.3 MgCl2.6H2O, 2 CaCl2.2H2O. Recordings were amplified using a Multiclamp 700B amplifier (Molecular Devices) and digitized using a Digidata 1440A (Molecular Devices). Patch electrodes (ranging from 6 to 9 MΩ) were filled with intracellular solution containing (in mM): 120 KMeSO4, 10 KCl, 10 HEPES, 8 NaCl, 4 Mg-ATP, 0.3 Na-GTP, and 0.3 Tris-base. The intracellular solution was also supplemented with 5 mM of biocytin to ensure cell morphological reconstruction.

### Histology

Animals were deeply anesthetized with tiletamine/zolazepam (Zoletil, 40 mg/kg) and medetomidine (Domitor, 0.4 mg/kg) and perfused transcardially with 4% paraformaldehyde. Serial frontal sections (100 μm) were performed using a vibrating microtome (Leica Biosystems) and mounted on glass slides. Images were taken using a fluorescence stereomicroscope (Olympus) and a wide-field microscope equipped with a structured illumination system (Zeiss). Measurements and tridimensional brain reconstructions were performed using Free-D.^14^

### Statistics

To study the age-related progression of seizure parameters (number, duration, amplitude, proportion of spikes), we computed the best-fit regression lines of these parameters against age, and considered the steepness of the regression lines as a descriptor of how epilepsy progresses with age. In order to compare epilepsy progression among groups, we compared slopes and intercepts of these regression lines, and estimated the probability that the elevations of these lines are different. To compare the effect sizes of excitability manipulation on the quantified endpoints (numbers of seizures and proportion of spikes within cortical discharges in 5-6-month-old rats), we computed the mean differences and their bootstrap 95% confidence intervals, and used Cumming estimation plots, as described by Ho *et al*.^15^ We performed statistical analyses with Prism 9 (GraphPad Software). Significance level was set at P < 0.05. We generated Cumming and Gardner-Altman estimation plots as described by Ho *et al*.^15^ and Violin SuperPlots as described by Kenny and Schoen.^16^

### Data availability

The data that support the findings of this study are available from the corresponding author, upon reasonable request.

## Results

### Adult bilateral Dcx-KD rats harbor bilateral SBH and exhibit chronic spontaneous seizures

We studied bilateral Dcx-KD rats as a genetic model of malformation of cortical development causing epilepsy. We generated cohorts of these rats using tripolar in utero electroporation, a method enabling targeted, bi-hemispheric genetic manipulations in the developing isocortex (Fig. 1A and Supplementary Fig. 1A, C). Adult bilateral Dcx-KD rats harbored prominent bilateral SBH extending rostrocaudally in the white matter and resembling that seen in humans (Fig. 1B-D), and exhibited chronic spontaneous non-convulsive seizures, as we reported earlier.^12^ These seizures in adulthood consisted of spike-and-wave discharges (SWDs) composed of trains of spikes (S) followed by associated slow waves (W), and referred to as spike-and-wave (SW) complexes (Fig. 1E1). No SWDs were detected, neither in sham-operated rats (injection only or tripolar electroporation only), nor in rats electroporated with ineffective shRNAs with mutations creating mismatches and harboring no SBH, thus confirming their epileptic nature, as we described earlier.^12^ We found that the amplitude and velocity of amplitude of SW could serve as a unique descriptor of their electrographic signature (Fig. 1E2), distinct to that of rhythmic waves or background noise (Fig. 1F). We thus developed an ad hoc seizure detection algorithm based on a character recognition approach utilizing these so-called phase portraits as alphabet characters (Supplementary Fig. 2A, B and Material and methods).

### Epilepsy progresses with age in juvenile rats with bilateral SBH

To investigate the development and potential age-related changes of epileptiform activity in bilateral Dcx-KD rats, we performed longitudinal electrocorticographic (ECoG) recordings over a three-month period with implanted telemetric devices equipped with motion sensors (Fig. 2A). We used our seizure detection algorithm to quantify seizures occurring over 24-hour periods, as well as their hourly counts, on a monthly basis from three to six months of age. We also quantified hourly motion counts, and examine their diurnal rhythmicity across the light-dark phase. Cosinor analysis of motion counts (Fig. 2B) confirmed that peaks of motion occurred in the dark phase at all ages, and that the rats showed significantly higher proportions of cumulated motion in the dark phase (median, 64.41% with 95% CI of median [60.68, 67.94], Fig. 2C), as expected from nocturnal animals. Hourly seizure counts however showed no clear rhythmicity and occurred in equal proportions in the light or dark phase (median, 48.82% with 95% CI of median [43.37, 50.41], Fig. 2C). Of note, the skewed distribution of inter-seizure intervals (Supplementary Fig 3) showed that seizures often occurred in clusters.

**Figure 2.**
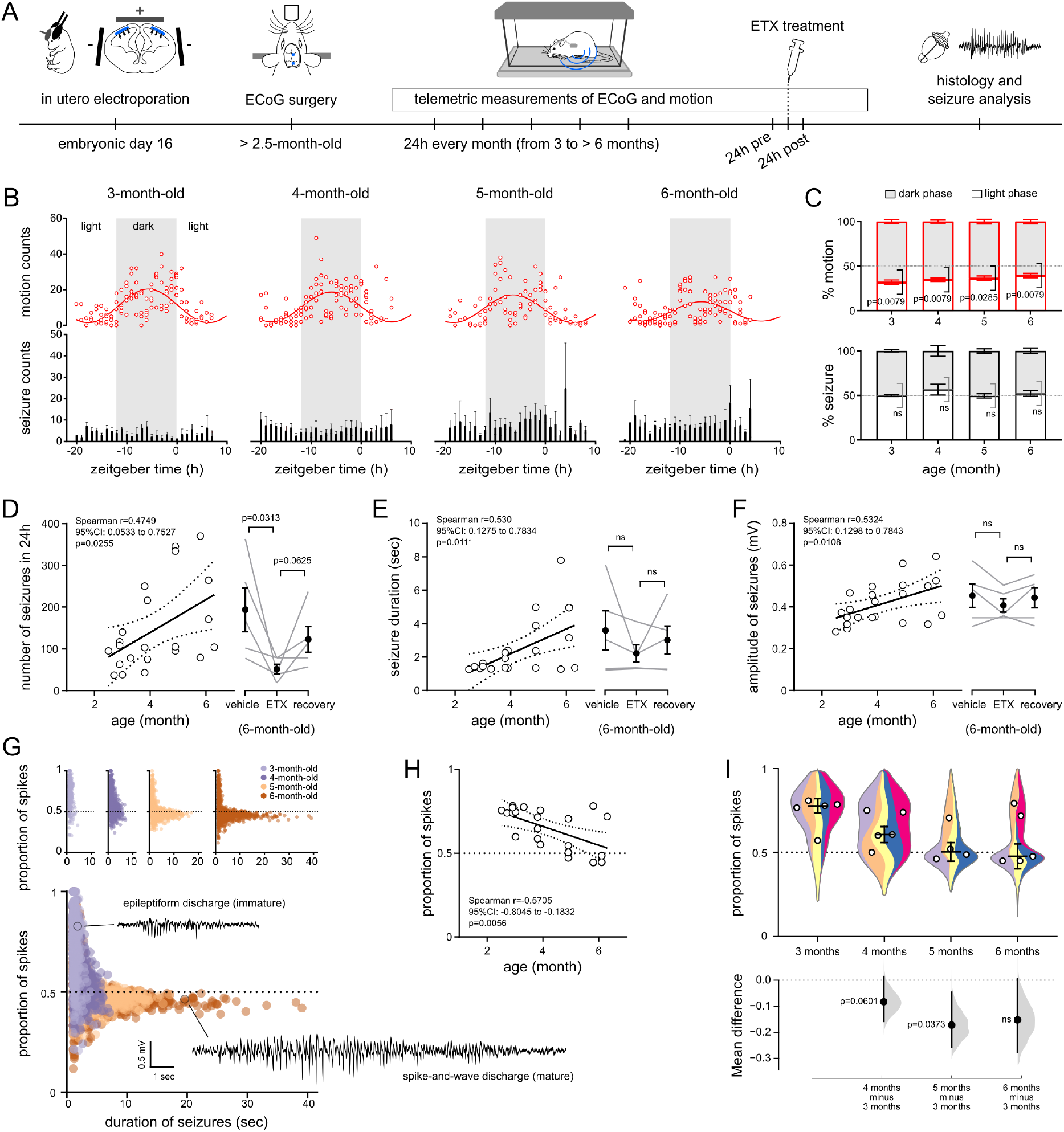
Age-dependent progression of epilepsy in bilateral SBH rats. **(A)** Experimental timeline. **(B)** Upper graphs show dot plots of hourly motion counts, with superimposed sine waves representing the Cosinor analysis of motion across the light-dark phase. Lower graphs show bar graphs of hourly seizure counts. Error bars show SEM. **(C)** Stacked bar graphs showing the proportion of motion (top) and seizure (bottom) occurring during the dark and light phase. Error bars show SEM. **(D-F)** Number of seizures in 24 hours (D, left), seizure duration (E, left) and amplitude (F, left) plotted as a function of age (in months), and best-fit linear regression lines. Open circles represent mean individual values, dotted lines show confidence intervals of the best-fit line. Line graphs showing the number of seizure in 24 hours (D, right), seizure duration (E, right) and amplitude (F, right) after vehicle and ethosuximide (ETX) administration, and during the recovery period after ETX administration (post-ETX) in the 6-month-old group. Black dots and error bars show mean values and SEM. **(G)** Proportion of spikes within cortical discharges plotted as a function of seizure duration in 4 age groups. The 4 graphs are superimposed in B to show the progression from brief epileptiform discharges mostly composed of spikes in young bilateral Dcx-KD rats, to long-lasting spike- and-wave discharges composed of an equal proportion of spikes and slow waves in older bilateral Dcx-KD rats. **(H)** Proportion of spikes within cortical discharges plotted as a function of age (in months), and best-fit linear regression line showing an age-dependent progression of discharges patterns in bilateral Dcx-KD rats. Open circles represent mean individual values, dotted lines show confidence intervals of the best-fit line. **(I)** The upper graph shows violin superplots (top) of the proportion of spikes within cortical discharges in 4 age groups. Colored stripes show the normalized density estimates of individual discharges for each rat. Outlines show the overall density estimate of the age group. Open circles show median values for each rat, and horizontal lines and error bars show mean and SEM for the age group. The lower graph shows a Cumming estimation plot illustrating the mean difference compared to the 3 month age group. Mean differences are plotted as bootstrap sampling distributions (light grey areas), black dots and vertical error bars show mean difference and 95% confidence intervals.

We next evaluated the correlation between the age of the rats and the number of seizures occurring over a 24h period, and with their duration or amplitude. We found significant positive relationships between the age and the number (Spearman *r* = 0.4749, *p* = 0.0255, Fig. 2D), the duration (Spearman *r* = 0.530, *p* = 0.0111, Fig. 2E) and the amplitude of seizures (Spearman *r* = 0.5324, *p* = 0.0108, Fig. 2F), suggesting an age-dependent evolution of seizures in bilateral Dcx-KD rats. Consistent with the pharmacological properties of SWDs in humans and rodent models, ethosuximide significantly reduced seizure numbers in six-month-old rats (*p* = 0.0313, Wilcoxon test, Fig. 2D), while the duration or amplitude of remaining seizures were unchanged (Fig. 2E, F). Last, we examined if the age-dependent changes in the number, duration and amplitude of seizures were associated with changes in their electrographic patterns. We quantified the proportion of spikes (vs. waves) within cortical discharges, plotted them as a function of seizure duration, and compared their distribution across ages. This analysis revealed that seizures gradually progresses from brief epileptiform discharges mostly composed of spikes in three-month-old bilateral Dcx-KD rats, to long-lasting fully developed SWDs composed of an equal proportion of spikes and waves from five months onward (Fig. 2G). We found a significant negative relationship between the age of the rats and the mean proportion of spikes within cortical discharges (Spearman *r* = -0.5705, *p* = 0.0056, Fig. 2H). We confirmed this observation by plotting both individual and pooled distributions of spike proportions per cortical discharges for individual rats and for the rats of the same age group (Fig. 2I). This revealed that, despite inter-individual differences, cortical discharges tended to comprise an equal proportion of spikes and waves by five months of age, and we considered that rats have reached a mature stage of SWDs.

Altogether, these observations indicate that epilepsy gradually progresses in juvenile rats with bilateral SBH, with prominent changes in the occurrence, duration and amplitude of seizures, and in the properties of their electrographic patterns.

### Early targeted suppression of neuronal excitability in SBH and not in the normotopic cortex modifies epileptogenesis

To dissect the contribution of SBH and that of the normotopic cortex (NCx) to the development and age-dependent progression of epilepsy, we utilized a targeted molecular genetic approach for suppressing neuronal excitability.^17^ We created cohorts of bilateral Dcx-KD rats functionally expressing either in SBH, or in the NCx, an inwardly rectifying potassium channel, Kir2.1, that we and others have previously used to manipulate excitability.^18,19,20^ We first confirmed with patch-clamp recordings that constitutive Kir2.1 expression in cortical neurons is associated with a significantly more hyperpolarized resting membrane potential than controls, consistent with a decreased excitability (mean difference, -18.36 mV, with 95% CI of mean [-24.49, -12.28], *p* = 0.00008, Mann-Whitney test, Supplementary Fig. 4).

We generated rats with SBH-targeted Kir2.1 expression (referred to as rats with silenced SBH in Fig. 3) by injection and tripolar co-electroporation of plasmids encoding GFP, Kir2.1 and Dcx shRNAs, respectively (Supplementary Fig. 5A1, A2). Of the total population of Kir2.1-expressing cells, those located in the SBH ranged between 37.39% and 98.34% with this experimental setting (Supplementary Fig. 6A). We then longitudinally monitored the age-related changes of epileptiform activity in rats with silenced SBH using telemetric ECoG recordings and the same experimental timeline as for un-manipulated bilateral Dcx-KD rats. We used the same parameters as described above to describe seizure progression. We calculated the best-fit regression lines and used them to describe the progression of seizure parameters with age, and to compare bilateral Dcx-KD rats with silenced SBH and un-manipulated ones. We found that rats with silenced SBH tended to have lower slope and elevation values for the number of seizures over a 24h period (elevation, *p* = 0.0580, Fig. 3A1), their duration (elevation, *p* = 0.0190, Fig. 3A2) and amplitude (elevation, *p* < 0.0001, Fig. 3A3), and for proportion of spikes within cortical discharges (elevation, *p* = 0.0123, Fig 3A4). Plots of the proportion of spikes as a function of seizure duration across ages also revealed a slower progression towards fully developed SWDs (Fig. 3A5). Because un-manipulated bilateral Dcx-KD rats harbor fully developed SWDs from five months onward, we compared the pooled numbers of seizures in five-six-month-old rats with silenced SBH to that of un-manipulated ones (Fig. 3C). We found that rats with silenced SBH had significantly lower numbers of seizures occurring over a 24h period than un-manipulated controls (mean difference, -94.77 seizures, with 95% CI of mean difference [-183, -9.61], *p* = 0.0360, Mann-Whitney test). Last, the proportion of spikes within cortical discharges tended to be higher than that of un-manipulated Dcx-KD rats, suggesting a lower maturation pace of SWDs (mean difference, 0.11, with 95% CI of mean difference [-0.01, 0.21], *p* = 0.0596, Mann-Whitney test, Fig. 3D). These findings are consistent with a modified epileptogenesis in bilateral Dcx-KD rats with silenced SBH, characterized by a slower age-related increase of seizure numbers and a slower progression towards fully developed SWDs.

**Figure 3.**
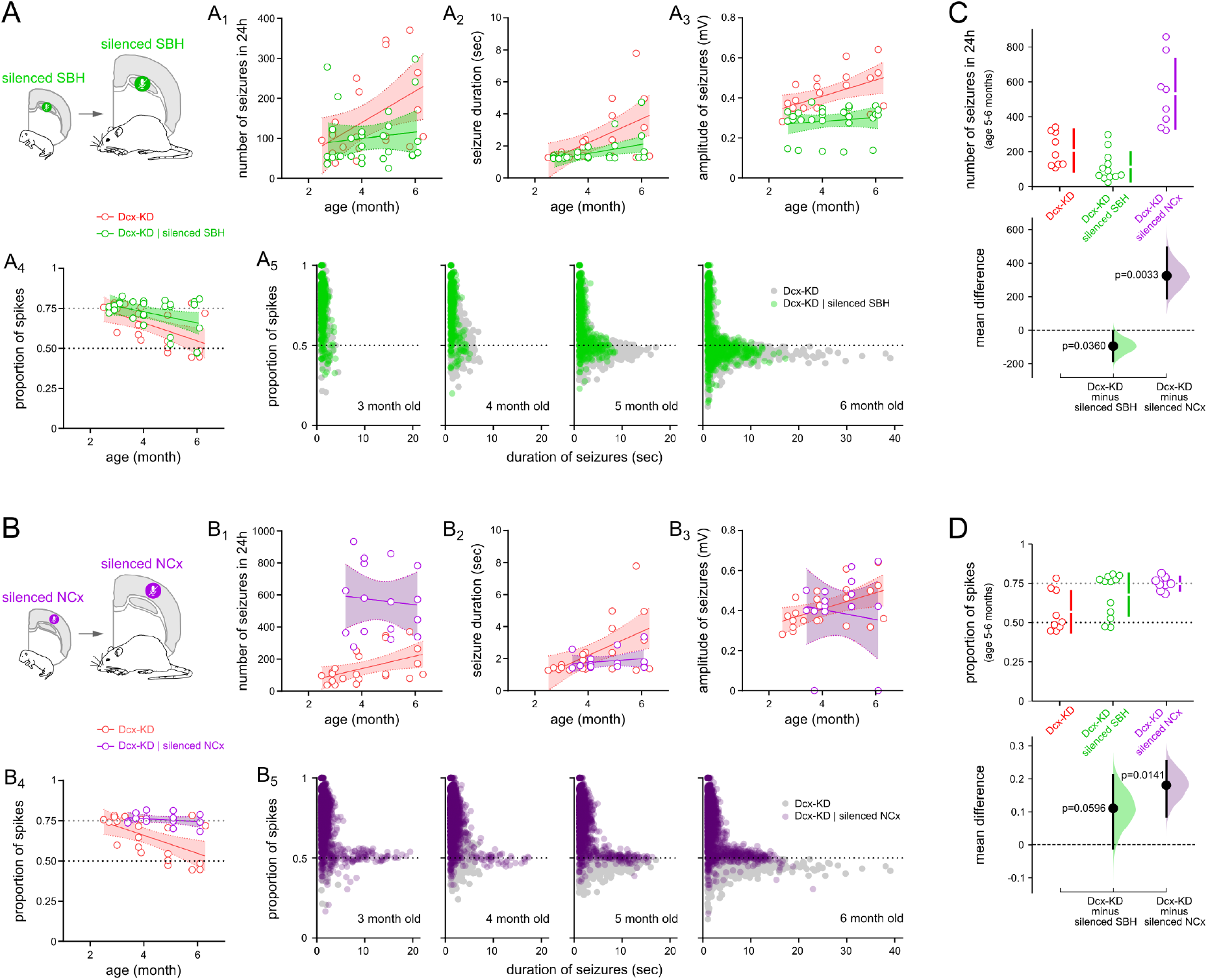
Early chronic suppression of neuronal excitability in the SBH of bilateral Dcx-KD rats modifies epileptogenesis, whereas early chronic suppression of neuronal excitability in the normotopic cortex worsens it. **(A, B)** Schematic views of the experimental design. **(A1-A4, B1-B4)** Number of seizures in 24 hours (A1, B1), seizure duration (A2, B2) and amplitude (A3, B3), and proportion of spikes within cortical discharges (A4, B4) plotted as a function of age (in months), and best-fit linear regression lines, in bilateral Dcx-KD rats (red) as compared to bilateral Dcx-KD rats with chronically suppressed neuronal excitability in SBH (silenced SBH, green) or in the normotopic cortex (silenced NCx, purple). Open circles represent mean individual values, shaded areas show confidence intervals of the best-fit lines. **(A5, B5)** Proportion of spikes within cortical discharges plotted as a function of seizure duration in 4 age groups. Grey circles correspond to bilateral Dcx-KD rats, green circles correspond to bilateral Dcx-KD rats with chronically silenced SBH, and purple circles to bilateral Dcx-KD rats with chronically silenced NCx. **(C, D)** Cumming estimation plots illustrating the mean difference in the number of seizures (C) and in the proportion of spikes within cortical discharges (D) in bilateral Dcx-KD rats compared to bilateral Dcx-KD rats with chronically silenced SBH (green) and NCx (purple). Individual values are plotted on upper graphs, with vertical bars illustrating mean and 95%CI. On lower graphs, mean differences are plotted as bootstrap sampling distributions (shaded areas). Black dots with vertical error bars show mean difference and 95%CI.

We next evaluated the effects of a similar manipulation in the normotopic cortex of bilateral Dcx-KD rats. To generate rats with NCx-targeted Kir2.1 expression (referred to as rats with silenced NCx in Fig. 3), we used tripolar in utero electroporation and a two-step procedure (Supplementary Fig. 1B). We first bilaterally injected and co-electroporated plasmids encoding mCherry and Kir2.1 to express these transgenes in the NCx; then, 30 minutes later, we bilaterally injected and co-electroporated plasmids encoding GFP and Dcx shRNAs to create bilateral SBH (Supplementary Fig. 5B1, B2). Of the total population of Kir2.1-expressing cells, those located in the NCx ranged between 67.24% and 94.42% with this experimental setting (Supplementary Fig. 6B). We then longitudinally monitored the age-related changes of epileptiform activity with the same experimental and analysis pipeline as described above. We found that rats with silenced NCx had strikingly high numbers of seizures from 3 months onward, with no clear age-related evolution, suggesting that they had already reached a plateau number of seizures (Fig. 3B1). The duration (Fig. 3B2) and amplitude (Fig. 3B3) of seizures remained similarly stable with age. Interestingly, the proportion of spikes within cortical discharges did not change with age, and remained in the range values of immature epileptiform discharges found in young un-manipulated Dcx-KD rats (Fig. 3B4, 3B5). We compared the pooled numbers of seizures in five-six-month-old rats with silenced NCx to that of un-manipulated ones and found that rats with silenced NCx had significantly higher numbers of seizures occurring over a 24h period (mean difference, 325.30 seizures, with 95% CI of mean difference [190.97, 493.90], *p* = 0.0033, Mann-Whitney test, Fig. 3C). Last, rats with silenced NCx had a significantly higher proportion of spikes within cortical discharges than that of un-manipulated Dcx-KD rats (mean difference, 0.18, with 95% CI of mean difference [0.08, 0.25], *p* = 0.0141, Mann-Whitney test, Fig. 3D). These findings are consistent with an exacerbated epileptogenesis in rats with silenced NCx, characterized by strongly increased numbers of epileptiform discharges that, however, remained immature and relatively shorter.

To confirm that the two-step in utero electroporation procedure did not by itself contribute to the exacerbated epileptogenesis, we generated rats with NCx-targeted mCherry expression. We first bilaterally injected and electroporated plasmids encoding mCherry; then, 30 minutes later, we bilaterally injected and co-electroporated plasmids encoding GFP and Dcx shRNAs to create bilateral SBH. We then longitudinally monitored the age-related changes of epileptiform activity with the same experimental and analysis pipeline as used before. We found that Dcx-KD rats with NCx-targeted mCherry expression did not significantly differ from unmanipulated Dcx-KD rats for all seizure parameters studied (Supplementary Fig. 7).

Together, these observations reveal that an early, targeted suppression of neuronal excitability in either SBH or the NCx of bilateral Dcx-KD rats has a strikingly distinct impact on epileptogenesis, leading to either a favorable improvement of epilepsy progression, or to its exacerbation.

### Suppression of neuronal excitability in the SBH or NCx of rats with active epilepsy is ineffective

We next investigated if suppressing neuronal excitability in the SBH or NCx could have a beneficial impact for seizures in Dcx-KD rats with active epilepsy. To gain a temporal control on the onset of excitability suppression, we used a molecular genetic approach enabling a conditional, 4-hydroxy-tamoxifen (4OHT)- and Cre-dependent expression of Kir2.1 channels (Supplementary Fig. 4, 5). With this approach, functional expression of Kir2.1 channels is induced with a single intraperitoneal injection of 4OHT (20 mg/kg), and is associated with significantly more hyperpolarized resting membrane potentials, as confirmed with patch-clamp recordings (mean difference, -13.40 mV, with 95% CI of mean [-17.31, -8.74], *p* = 0.0005, Mann-Whitney test, Supplementary Fig. 4).

We generated rats with conditional SBH-targeted Kir2.1 expression (referred to as rats with silenced SBH in Fig. 4A) using the same experimental approach as for the early silencing, but with different plasmid conditions (see Supplementary Fig. 5C1, C2). To generate rats with conditional NCx-targeted Kir2.1 expression (referred to as rats with silenced NCx in Fig. 4B), we used two-step tripolar in utero electroporation and the plasmid conditions described in Supplementary Fig. 5D1, D2.

**Figure 4.**
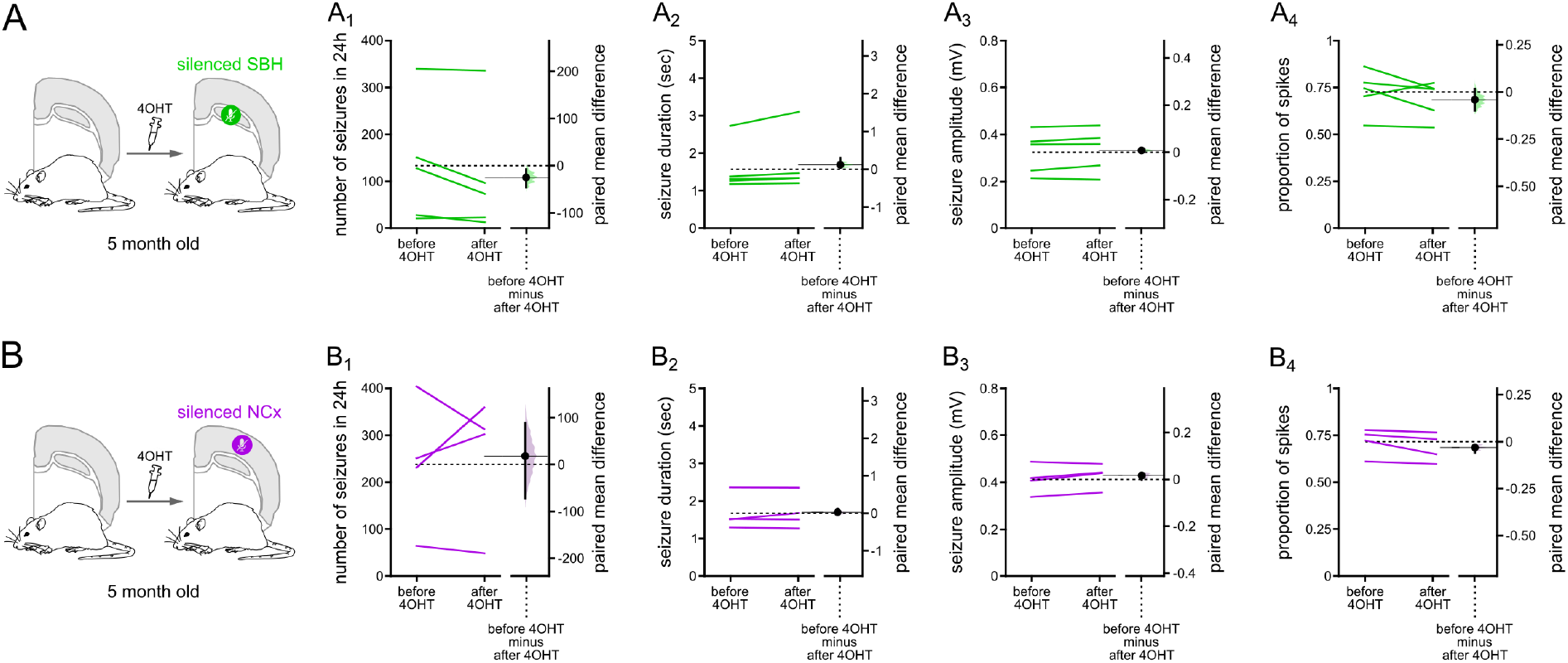
Suppression of neuronal excitability in either the SBH or the normotopic cortex is ineffective for ameliorating seizure phenotype in 5-month-old bilateral Dcx-KD rats with active epilepsy. **(A, B)** Schematic views of the experimental design. **(A1-A4, B1-B4)** Gardner-Altman estimation plots of the paired mean difference in the number of seizures in 24h (A1, B1), in the duration (A2, B2) and amplitude (A3, B3) of seizures, and in the proportion of spikes within cortical discharges (A4, B4) before and after suppression of neuronal excitability in the SBH (A1-A4) or in the NCx (B1-B4) of bilateral Dcx-KD rats, as induced by 4-hydroxy-tamoxifen (4OHT) injection. Each paired set of observations is connected by a line. The paired mean difference is plotted on a floating axis (right) as a bootstrap sampling distribution. Black dots with vertical error bars show mean difference and 95%CI.

Because bilateral Dcx-KD rats harbor fully developed SWDs from five months onward (see Fig. 2), we injected 4OHT in five-month-old rats. We then examined the short-term impact of the suppression of neuronal excitability by comparing seizure parameters one day before, and six days after 4OHT-induced Kir2.1 expression. In the two groups of rats, with either silenced SBH or NCx, we found no significant change in the numbers (Fig. 4A1, B1), duration (Fig. 4A2, B2) and amplitude of seizures (Fig. 4A3, B3), or in the proportion of spikes within cortical discharges (Fig. 4A4, B4). Together, these observations suggest that the approach has no beneficial impact in bilateral Dcx-KD rats with fully established epilepsy.

## Discussion

The pathological substrate and dynamic changes leading to the development and progression of epilepsy remain largely elusive in malformations of cortical development. In the present study, we have characterized the development and age-related progression of epilepsy in a rat model of bilateral SBH with chronic spontaneous seizures. We report a gradual evolution of epilepsy with age, with prominent quantitative and qualitative age-related changes in seizure properties and patterns, accompanying the progression towards a fully developed pattern of SWD seen in adulthood. We further show that early and targeted manipulations of neuronal excitability in either the SBH or the overlying cortex have drastic effects on this age-related epilepsy progression. While an early suppression of SBH excitability has clear antiepileptogenic effects, with a slower age-related increase of seizure numbers and a slower progression towards fully developed SWDs, the same manipulation in the normotopic cortex paradoxically exacerbates epileptogenesis. Last, we report that manipulating neuronal excitability with the same approach once epilepsy is established is ineffective, suggesting the existence of protracted critical periods for intervention.

Epilepsies are progressive brain network disorders, and especially those arising in childhood as a consequence of disrupted cortical development and aberrant circuit formation. Brains with SBH linked to DCX mutations in humans are mosaic,^21^ either caused by somatic mosaicism in male patients^22^ or related to X-inactivation in female patients.^9^ Accordingly, epilepsy in brains with SBH emerges and progresses in a context of ongoing cortical development, where both genetically normal and genetically altered neuronal populations interact together to form cortical networks. Hence, developing epileptogenic networks receive dual detrimental influences: those occurring as a direct consequence of the gene mutation, and those, likely reactive and non-genetic, arising indirectly or secondarily. In line with this, we have previously described in SBH rats both pathological alterations in the SBH,^23,24^ and secondary alterations in the overlying cortex,^20,23^ several weeks before epilepsy onset. Further, we have reported developmental circuit changes in the normotopic cortex that were distinct depending on SBH location: decreased circuit excitability and connectivity strength in the cortex located above SBH, increased excitability and connectivity in the cortex lateral to SBH.^25^ Such a complex combination comprising both local and distant changes suggests that brains with SBH harbor widespread circuit-level defects extending beyond the macroscopically visible heterotopia. This is in line with neuroimaging studies in patients with various types of MCDs, revealing widespread structural and functional circuit reorganization in focal cortical dysplasia (FCD),^26^ in periventricular nodular heterotopia (PVNH)^27^ and in SBH.^28^

In the present study, early manipulation of neuronal excitability has led to strikingly different consequences depending on the targeted area. These differences likely reflect distinct pathological mechanisms contributing to epileptogenesis in SBH – in which the causative mutation is the main contributor, and in the overlying cortex – in which secondary circuit-level reorganizations are prevailing as we demonstrated in Plantier *et al*.^25^ If both regions do concur to creating seizure-prone circuits, the weight of the SBH (and that of the causative gene) appears greater than that of the overlying cortex, as illustrated by a more favorable outcome of SBH-targeting manipulation. Although this remains to be tested, attenuating influences of the SBH by suppressing its excitability has likely prevented or attenuated both the pathogenic changes directly related to the causative gene, as well as the development and/or expansion of secondary circuit-level alterations. It would be equally important to clarify the possible contribution of altered patterns of neuronal firing in the SBH, and modified strength of functional network connectivity, to the ameliorated seizure outcomes. On the contrary, attenuating influences of the normotopic cortex was found insufficient, and, paradoxically, detrimental. This is not entirely unexpected, given that complex maladaptive circuit changes, combining both increased and decreased circuit excitability and connectivity, were found in the overlying cortex.^25^ In this context, the destabilizing effects of local excitability changes may have led to an exacerbation of cortical imbalance in the normotopic cortex, further precipitating epileptogenesis. Whether larger numbers of cells with manipulated neuronal excitability in the normotopic cortex would lead to greater cortical imbalance and even more severe seizure phenotypes remains to be investigated.

Timing of intervention matters, and excitability suppression in rats with fully established epilepsy was found inefficient. Although speculative, we propose several explanations for this. First, with a continuing progression of network dysfunction as epilepsy progresses, the spatial extend of circuit alterations is likely far greater than that of the targeted circuits, precluding any beneficial impact. Second, the targeted circuits may potentially fall outside of the most epileptogenic regions, and their precise location and properties require clarification. Last, circuit-level alterations whose development and/or expansion can be limited with early interventions are unlikely to regress once development has ended. These hypotheses are in agreement with neuroimaging studies suggesting that the strength of the abnormal circuitry increases over time, as epilepsy progresses, in patients with gray matter heterotopia.^27^ Therefore, the present work supports the notion that early corrective interventions, including surgical removal prior to drug-resistance, as recently proposed,^29^ may reduce morbidity, and improve neurocognitive outcomes.

Whether the favorably modified epileptogenesis described here has other beneficial effects remains to be investigated. The age of epilepsy onset in patients with SBH usually correlates with the severity of epilepsy and that of associated co-morbidities. Earlier ages of epilepsy onset, and larger proportions of drug-resistance, are observed in patients presenting with more severe intellectual and behavioral impairments, and with wider anatomical extend of SBH.^9,10^ We have previously reported that rats with unilateral SBH, and older ages of epilepsy onset, showed no cognitive impairments when tested prior to epilepsy onset.^30^ Whether any cognitive alterations are associated with epilepsy progression, or are already present before epilepsy onset in rats with bilateral SBH would require further work.

In a large series of patients described by Bahi-Buisson *et al*.^9^ 85% of SBH patients had seizure disorders, with epilepsy starting in infancy (35% of cases) or during childhood (45% of cases), and often associated with an evolution of seizure types or electrographic patterns with age. Here, we have described in SBH rats an age-related progression of electrographic patterns where epileptiform discharges finally shaped into fully developed SWDs. Longitudinal EEG monitoring of individual animals, as we have done here, is rarely performed over prolonged periods in rodent models of MCDs. Kao *et al*.^31^ recently studied epileptogenesis in a rat model of Depdc5-related FCD type II, and documented a similar progression of immature electrographic patterns towards mature forms of cortical discharges. Using both intermittent and continuous EEG monitoring, they found that mature patterns gradually emerge from brief runs of SW complexes that progressively last longer, and finally morph into well-formed SWDs.^31^ Likewise, in the GAERS rat model of absence-type epilepsy, epileptiform activity gradually progresses during epileptogenesis. Oscillatory epileptiform discharges emerge first, become progressively intermingled with SW complexes, and finally shape into fully mature SWDs.^32^ In GAERS rats, this progression towards mature discharges is accompanied by progressively increased excitability of cortical neurons and strengthening of local synaptic activity,^32^ increased structural connectivity in the cortex,^33^ and altered developmental sequences of cortical wiring.^34^ In bilateral SBH rats, the evolution of electrographic patterns likely engage similar mechanisms, and combines intrinsic and circuit excitability changes, modified excitation-inhibition ratio, and altered cortical connectivity, as we demonstrated previously.^25^ Because we have observed a slower progression towards fully mature SWDs upon manipulation of excitability of both the SBH, and the normotopic cortex, this suggests that the two regions contribute to the maturation of electrographic patterns. Interestingly, reciprocal connections between the SBH and the normotopic cortex were described in SBH patients,^28^ in PVNH patients,^27,35,36^ and in murine models of SBH and PVNH (Vermoyal, Hardy *et al*., unpublished work) in keeping with this idea. Whether these reciprocal connections do contribute to the propagation or to the generalization of seizures in bilateral SBH rats would require further in vivo investigations with multi-site and multi-electrode recordings with precise targeting of the normotopic cortex and SBH.

To conclude, our longitudinal monitoring of epilepsy progression in rats with bilateral SBH reveals a gradual evolution of seizure properties and patterns during the course of epileptogenesis, before finally reaching a fully developed discharge pattern. Further, through timed and spatially targeted manipulation of neuronal excitability, we demonstrate that a complex combination of developmental alterations in both the SBH and the cortex contributes to epileptogenesis. Importantly, our work also suggests that early corrective interventions may help attenuating the development or expansion of circuit-level alterations, with a positive impact on altered developmental trajectories. Because epileptogenesis and its treatment are priorities in epilepsy research and care,^2^ our work may contribute to a better understanding of the pathological substrate leading to epileptogenesis, and to the development of novel corrective interventions.

## Abbreviations

ECoG: ElectroCorticoGram
DCX: doublecortin
Dcx-KD: doublecortin knockdown
FCD: focal cortical dysplasia
MCD: malformations of cortical development
PVNH: periventricular nodular heterotopia
SBH: subcortical band heterotopia
NCx: normotopic cortex
shRNA: short hairpin RNA
SW: spike-and-wave
SWD: spike-and-wave discharge

## Acknowledgements

We thank Dr L. Cancedda for providing a prototype of the third electrode used for tripolar electroporation at the initial stage of the study, Dr J. LoTurco for Dcx shRNAs plasmids and Dr Y. Tagawa for Kir2.1 plasmids. We also thank the animal facility (PPGI, INMED, Marseille), the imaging facility (INMAGIC, INMED, Marseille), and the molecular and cellular biology facility (PBMC, INMED, Marseille).

## Supplementary material

Supplementary material is available online.

## Supplementary Figures

**Supplementary Figure 1.**
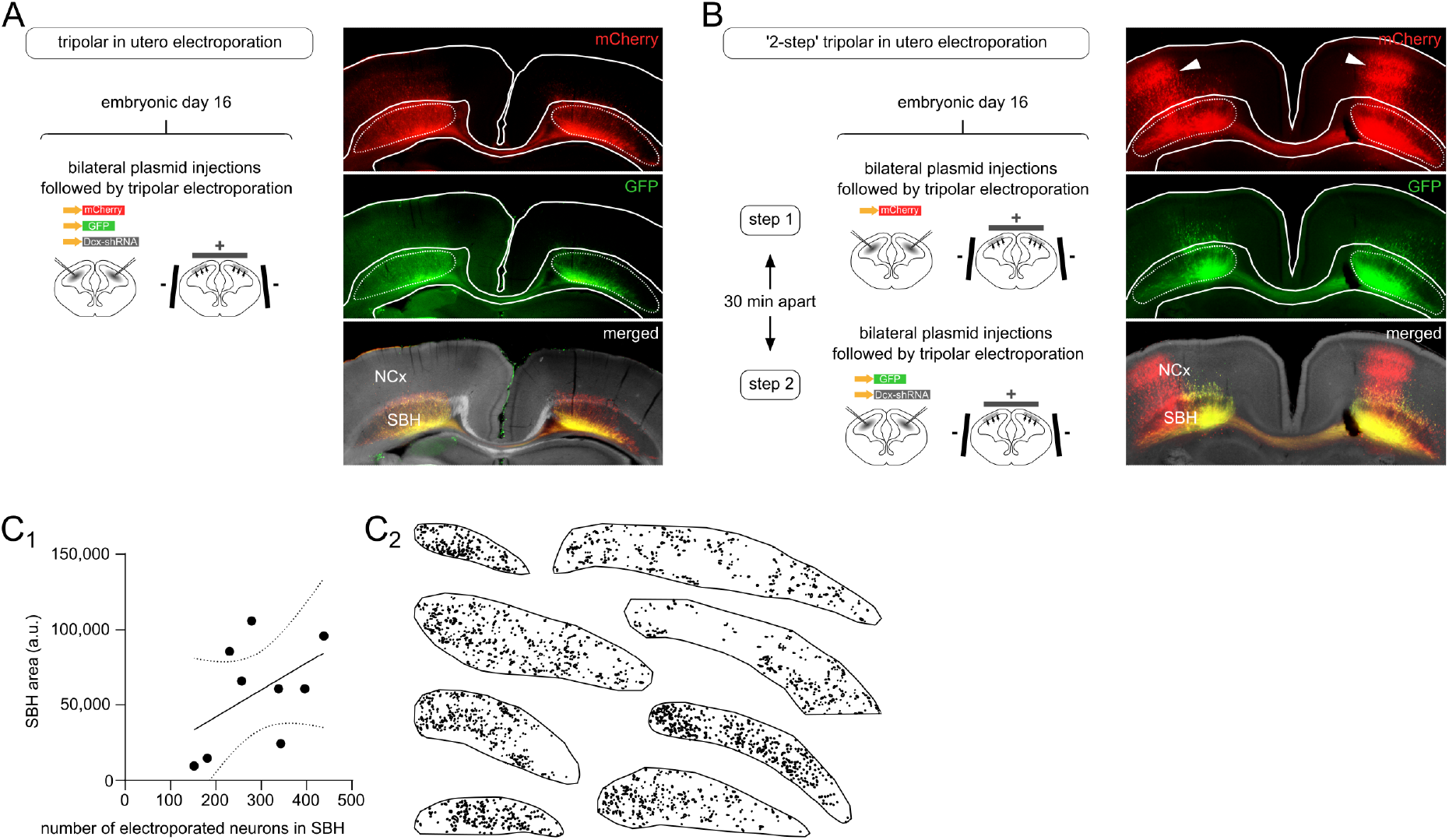
Tripolar in utero electroporation settings utilized to generate bilateral Dcx-KD rats. **(A)** (left) Schematic view of the tripolar in utero electroporation settings for bilateral electroporation. Lateral ventricles of embryonic day 16 embryos are bilaterally injected with plasmids encoding mCherry (0.5 μg/μl), GFP (0.5 μg/μl) and Dcx shRNAs (1 μg/μl). Electroporation is accomplished by delivering voltage pulses across 3 electrodes positioned bilaterally and at the brain midline. (right) Representative example of a postnatal day 7 brain with bilateral subcortical band heterotopia (SBH). Bilateral SBH composed of neurons co-expressing mCherry and GFP are located in the white matter below the normotopic cortex (NCx). **(B)** (left) Schematic view of the tripolar in utero electroporation settings for 2-step bilateral electroporation. Embryonic day 16 embryos are bilaterally injected and electroporated with mCherry plasmids (0.5 μg/μl), 30 minutes before plasmids encoding GFP (0.5 μg/μl) and Dcx shRNAs (1 μg/μl). Expression of mCherry and GFP is driven by the synthetic cytomegalovirus enhancer chicken beta-actin hybrid promoter (CAG promoter), enabling strong and ubiquitous expression (Niwa et al., Gene 1991). Expression of Dcx shRNAs is driven by the mouse U6 RNA polymerase III promoter (Yu et al., PNAS 2002), enabling effective knockdown of Dcx expression in vivo (Bai et al., Nat Neurosci 2003). (right) Representative example of a postnatal day 7 brain with a GFP-expressing bilateral subcortical band heterotopia (SBH) and a mCherry-expressing normotopic cortex (NCx). **(C)** (C1) Area of the SBH plotted as a function of the number of electroporated neurons in SBH, and best-fit linear regression lines illustrating a linear relationship. (C2) representative examples of reconstructed SBH with superimposed electroporated neurons (black dots).

**Supplementary Figure 2.**
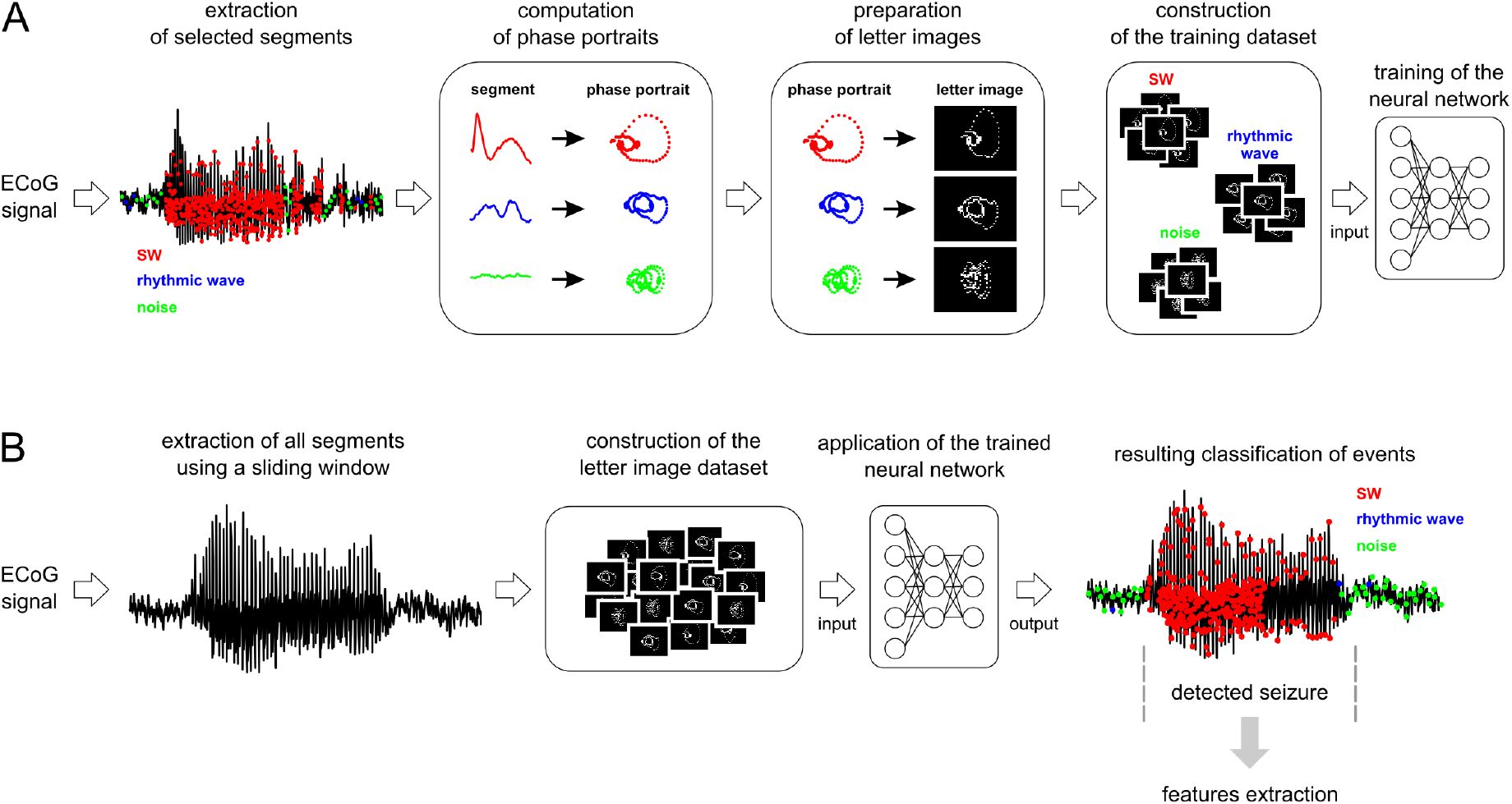
Seizure detection framework. **(A)** Seizure detection framework at the training phase. **(B)** Seizure detection framework at the application phase. See Material and methods for details.

**Supplementary Figure 3.**
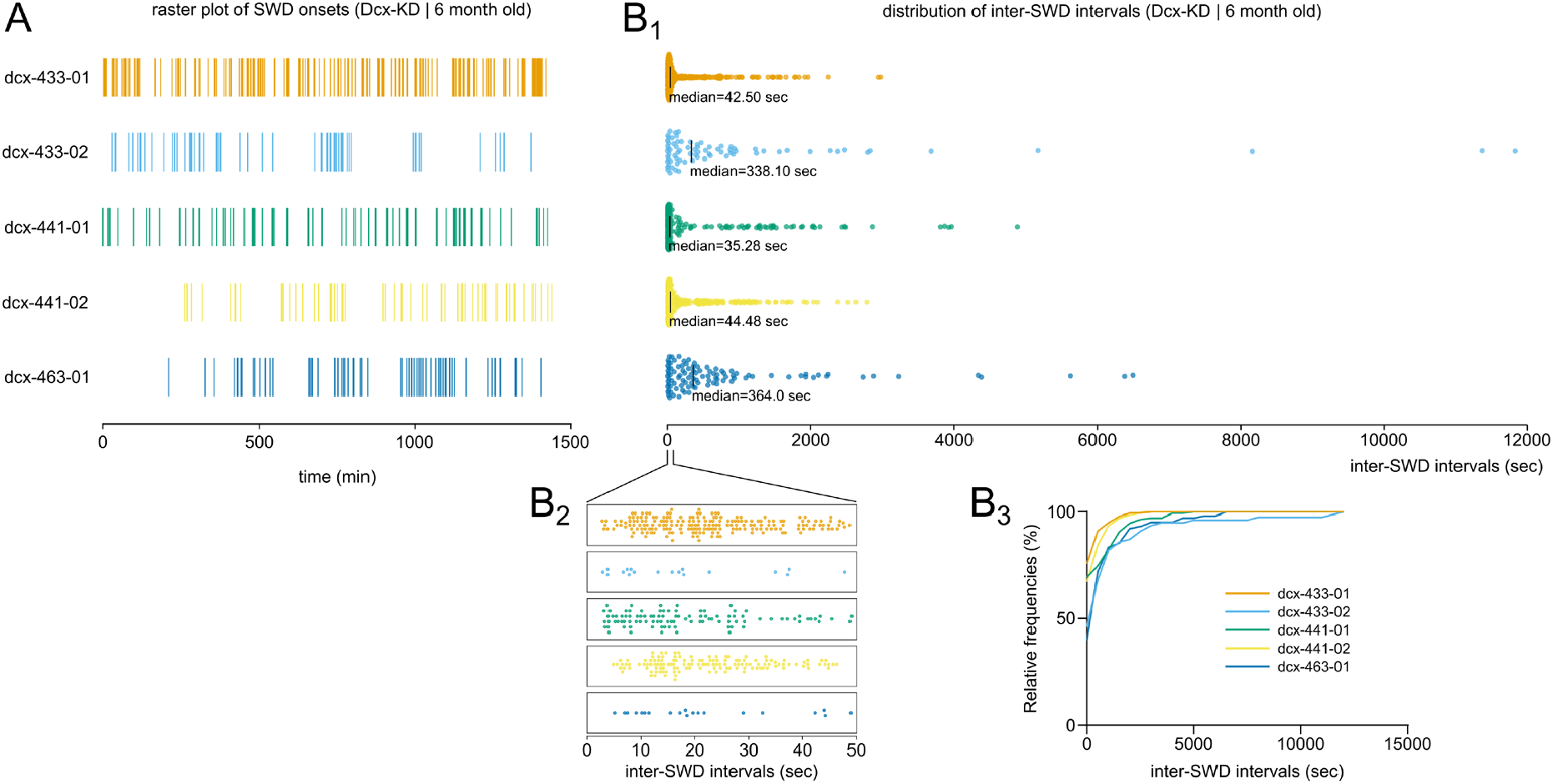
Inter-seizure intervals in bilateral Dcx-KD rats. (A) Raster plots of seizure (SWD) onsets in 5 bilateral Dcx-KD rats across a 24h recording period. (B1, B2) Dot plots of inter-SWD intervals in the same rats, with inter-SWD intervals shorter than 50 seconds viewed at an expanded timescale in B2. (B3) Cumulative frequency distribution of inter-SWD intervals in the same rats.

**Supplementary Figure 4.**
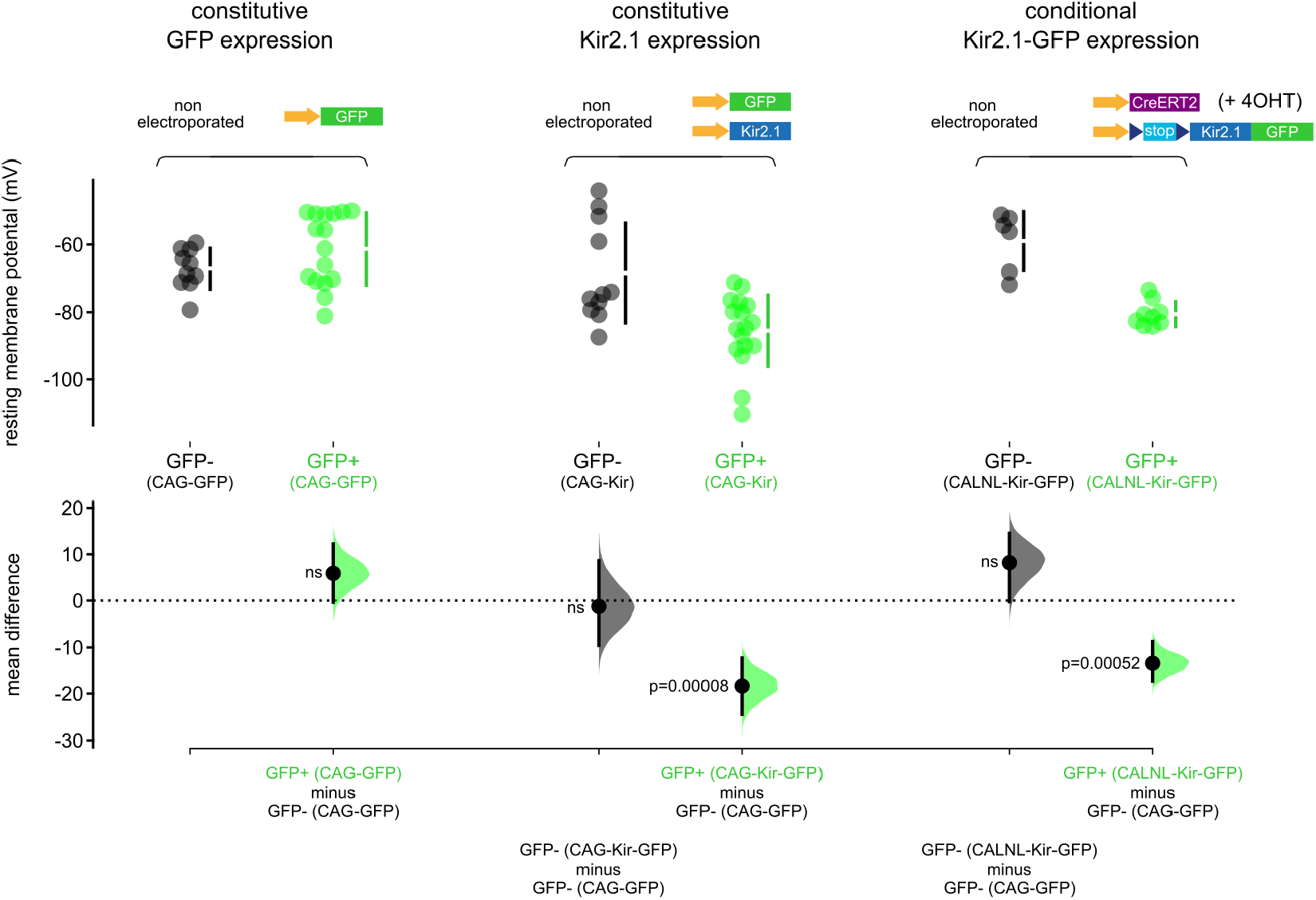
Neurons expressing Kir2.1 display a significantly more hyperpolarized resting membrane potential than controls, consistent with a decreased neuronal excitability. Cumming estimation plot illustrating the mean difference in the resting membrane potential of neurons constitutively expressing GFP (left, green) or Kir2.1 channels (middle, green), and of neurons conditionally expressing Kir2.1 channels after 4-hydroxy-tamoxifen (4OHT)-dependent and Cre-mediated recombination (right, green), as compared to non-electroporated controls (left, black). Individual values are plotted on upper graphs, with vertical bars illustrating mean and 95%CI. On lower graphs, mean differences are plotted as bootstrap sampling distributions (shaded areas). Black dots with vertical error bars show mean difference and 95%CI. Illustrated statistics correspond to those comparing the resting membrane potentials in GFP positive neurons (GFP+) from every conditions to that of the non-electroporated (GFP-) neurons in the CAG-GFP condition (left, black). Resting membrane potentials of non-electroporated neurons in all conditions show no significant difference, as well as that of neurons expressing GFP alone (these statistics are not illustrated here). Plasmid concentrations used: GFP (0.5 μg/μl), Kir2.1 (1 μg/μl), CreERT2 (1 μg/μl), stop-Kir2.1-GFP (1 μg/μl). Expression of all transgenes is driven by CAG promoters.

**Supplementary Figure 5.**
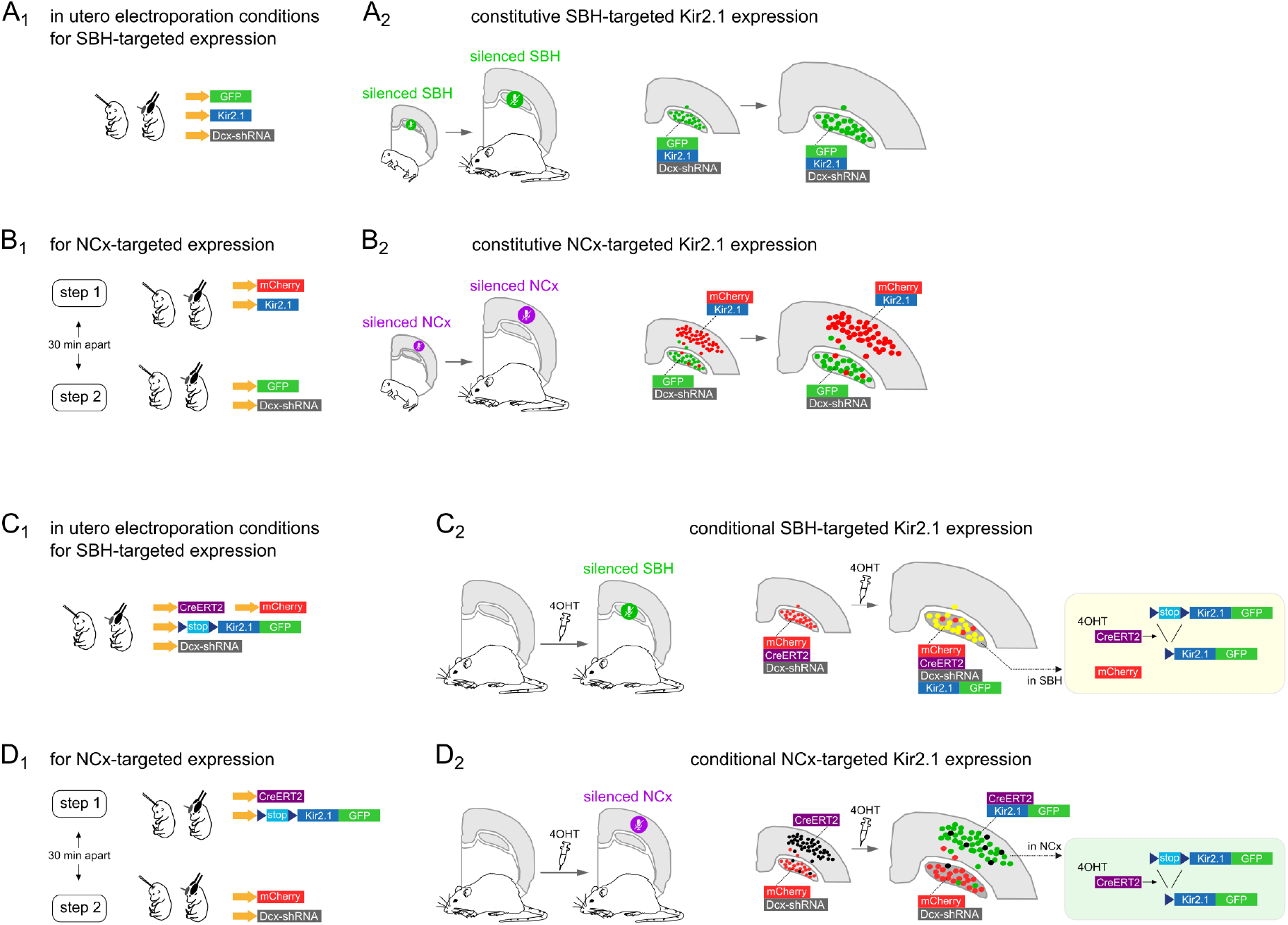
In utero electroporation conditions utilized for SBH- and NCx-targeted constitutive or conditional manipulation of neuronal excitability. **(A1, B1, C1, D1)** Schematic views of the plasmids and tripolar in utero electroporation settings utilized for SBH- (A1, C1) and NCx- (B1, D1) targeted expression. **(A2, B2, C2, D2)** Schematic views of the strategies for constitutive and conditional silencing of the SBH (A2, C2) and the normotopic cortex (B2, D2). For conditional approaches, a 4-hydroxy-tamoxifen (4OHT) activatable form of Cre recombinase is used to excise a floxed ‘stop’ cassette and to induce expression of Kir2.1 channels upon 4OHT injection (20 mg/kg). Plasmid concentrations used: GFP (0.5 μg/μl), mCherry (0.5 μg/μl), Kir2.1 (1 μg/μl), CreERT2 (1 μg/μl), stop-Kir2.1-GFP (1 μg/μl), Dcx-shRNA (1 μg/μl). Expression of all transgenes is driven by CAG promoters. Expression of Dcx shRNAs is driven by a U6 promoter.

**Supplementary Figure 6.**
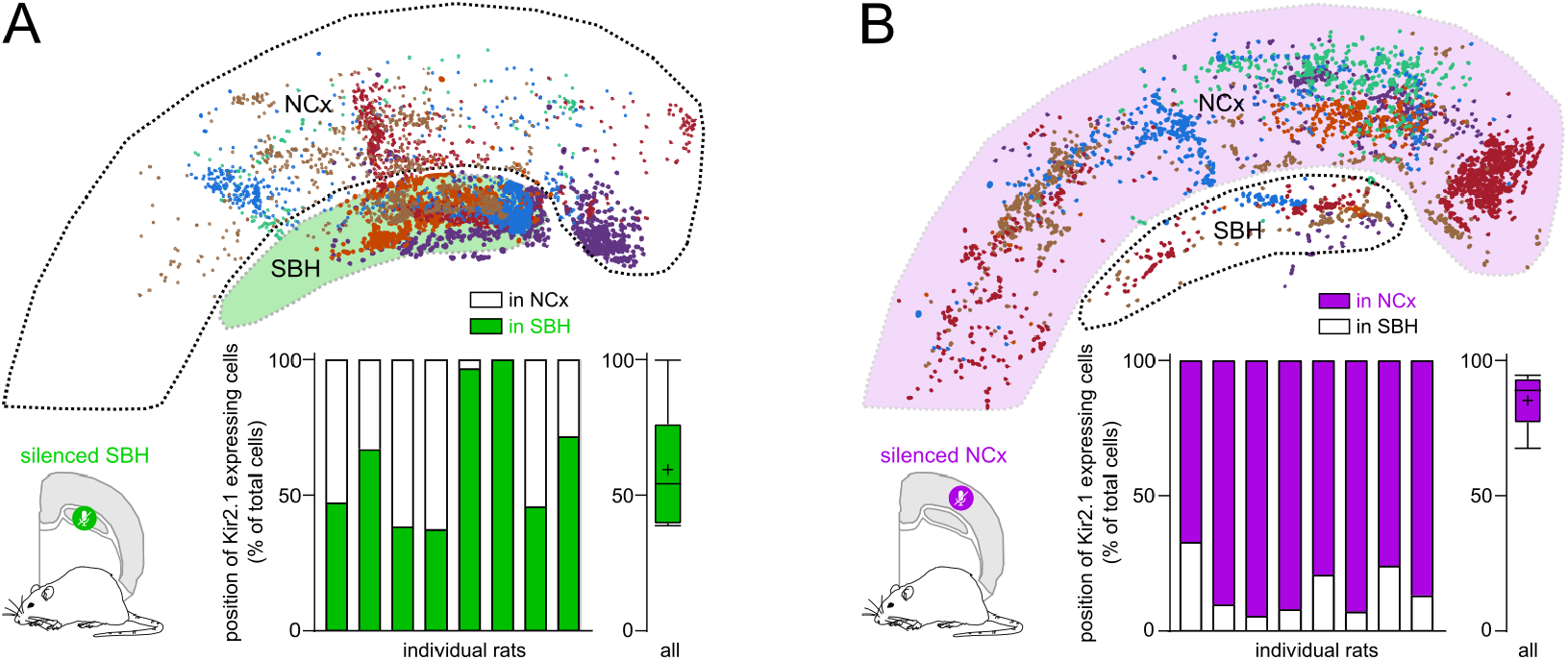
Distribution of Kir2.1-expressing cells in the SBH and normotopic cortex, in rats with SBH-targeted and in rats with NCx-targeted Kir2.1 expression. (top) Diagrams illustrating the distribution of Kir2.1-expressing cells (dots) in the normotopic cortex (NCx) and SBH of 8 individual rats with SBH-targeted (A) and NCx-targeted (B) Kir2.1 expression. Individual distributions are distinctly color-coded, realigned and superimposed onto a standardized brain section including NCx and SBH. (bottom) Stacked bar graphs illustrating the proportion of Kir2.1-expressing cells in the Ncx and SBH of 8 individual rats with SBH-targeted (A) and NCx-targeted (B) Kir2.1 expression. Box and whisker plots on the right show min-max values (whiskers), lower and upper quartiles (box), medians (horizontal line) and means (cross).

**Supplementary Figure 7.**
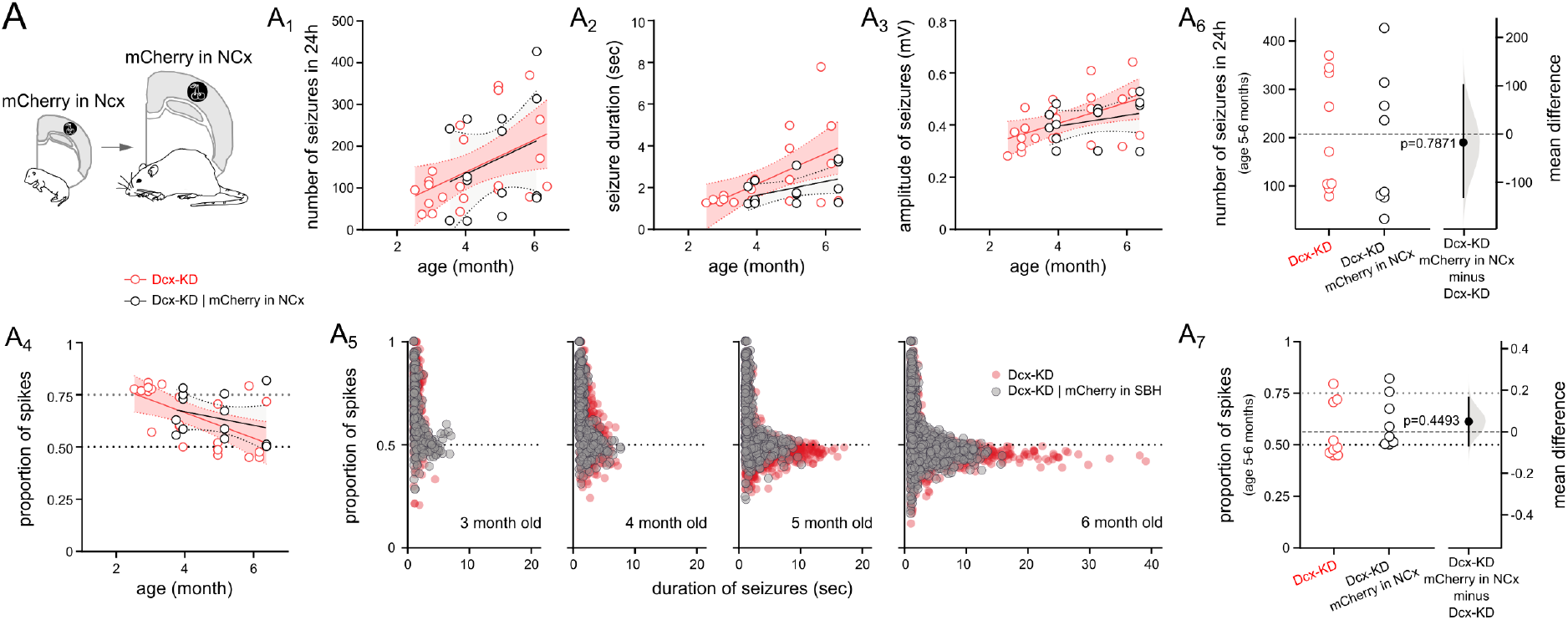
Seizure progression in rats with NCx-targeted mCherry expression did not significantly differ from that of unmanipulated Dcx-KD rats. (A) Schematic view of the experimental design. (A1-A4) Number of seizures in 24 hours (A1), seizure duration (A2) and amplitude (A3), and proportion of spikes within cortical discharges (A4) plotted as a function of age (in months), and best-fit linear regression lines, in bilateral Dcx-KD rats (red) as compared to bilateral Dcx-KD rats with NCx-targeted mCherry expression (mCherry in NCx, black). Open circles represent mean individual values, shaded areas show confidence intervals of the best-fit lines. (A5) Proportion of spikes within cortical discharges plotted as a function of seizure duration in 4 age groups. Red circles correspond to bilateral Dcx-KD rats, grey circles correspond to bilateral Dcx-KD rats with NCx-targeted mCherry expression. (A6, A7) Cumming estimation plots illustrating the mean difference in the number of seizures (A6) and in the proportion of spikes within cortical discharges (A7) in bilateral Dcx-KD rats (red) compared to bilateral Dcx-KD rats with NCx-targeted mCherry expression (black). Individual values are plotted on the left axes, with vertical bars illustrating mean and 95%CI. On the right axes, mean differences are plotted as bootstrap sampling distributions (shaded areas). Black dots with vertical error bars show mean difference and 95%CI.

